# Trimethylamine N-oxide (TMAO) enhances substrate mechanical stability probed by single molecule magnetic tweezers

**DOI:** 10.1101/2022.04.26.489563

**Authors:** Deep Chaudhuri, Debojyoti Chowdhury, Soham Chakraborty, Akashdeep Dutta, Ayush Mistry, Shubhasis Haldar

**Affiliations:** Department of Chemistry, Ashoka University, Sonepat, Haryana, India; Department of Biological Sciences, Ashoka University, Sonepat, Haryana, India

## Abstract

Trimethylamine N-oxide (TMAO) is a well-known osmolyte to stabilize the folded proteins through a variety of mechanisms. Since mechanical strength of proteins is a critical determinant of its stabilization, TMAO might play a relevant role by favoring its folding dynamics or by enhancing its mechanical stability. To address this question, we have performed the single-molecule magnetic tweezers experiment to explore the TMAO effect on two structurally distinct substrates-protein L and talin. We observed that TMAO increases the mechanical stability of these proteins through increasing their unfolding force. Additionally, we are able to demonstrate that TMAO retards the unfolding kinetics, while accelerating the refolding kinetics under force; which eventually tilts the energy landscape towards the folded state. Interestingly, this TMAO-enhanced protein folding generates mechanical work output upto ∼67 zJ, allowing the protein folding under higher force regime. Overall this TMAO-enhanced mechanical stability provides a significant implication to folding-induced structural stability of proteins.

## INTRODUCTION

Many aquatic animals employ different small osmolytes to stabilize their intracellular proteins against urea-induced denaturation^1–3^. These protective molecules enhance thermodynamic stability of different proteins, signifying themselves as ‘chemical chaperones^2^. Especially, Trimethylamine N-oxide (TMAO) is a well-known osmolyte to prevent destabilizing effects of urea and other salts^4,5^. Additionally, TMAO has been reported to accelerate the collapsed and aggregated states of elastin-like polypeptide and amyloid β-peptide^3,6^. Recently, experimental and computational studies have revealed diverse mechanisms of TMAO-mediated protein stabilization, including depletion effects through its unfavourable interaction with protein backbone and its surfactant-like effect^3,7–11^. An optical tweezers-based force study has also suggested that TMAO modulates early folding events of T4 lysozyme, stabilizing its on-pathway intermediate^12^. Since mechanical stability of proteins is an indispensable aspect to understand their folding phenomena, it is plausible that TMAO could affect the protein folding by modulating their stability under force. However, despite well-established studies on how TMAO influences protein folding, the precise mechanism responsible for substrate mechanical stability remains elusive.

To address this question, we have performed single-molecule magnetic tweezers spectroscopy, which employs both the force-ramp and force-clamp methodology^13,14^. Using the force-ramp methodology, the applied force can be increased or decreased at a constant loading rate, detecting unfolding and refolding events at different forces on a single molecule; while the force-clamp methodology at a constant applied force allow us to measure the TMAO effect on both the folding dynamics and kinetics under equilibrium condition. Finally, the advantage of a larger force range of 0-120 pN allow us to observe the TMAO interaction with both the folded and unfolded states of the substrate proteins independently.

Here, we have investigated the TMAO effect on protein L as a model substrate, which has previously been used in force spectroscopic studies^14–17^. Since protein L is not biologically relevant to function under mechanical tension, we selected another substrate talin, which is well-known to work under physiological force and extensively used in force spectroscopy studies^13,18–20^. Interestingly, these two substrates are structurally different: protein L is predominantly a β-sheet containing substrate, whereas talin is an α-helical protein. Our results showed that TMAO mechanically stabilizes their substrates by increasing their unfolding force from ∼43 pN to ∼56 pN. We have also reported that TMAO enhances this mechanical stability through tilting the energy landscape towards the folded state, which has been reconciled from the increased folding probability data of the substrate proteins. Additionally, TMAO has been found to retard the unfolding kinetics and accelerate the refolding kinetics of protein L. Lastly, TMAO boost the mechanical work output of protein L folding upto two folds through assisting their folding at higher force. Overall this TMAO-enhanced mechanical stability represents a crucial aspect of folding-induced structural stability of proteins.

## Results

### Folding dynamics of protein L measured by single-molecule magnetic tweezers

Single-molecule magnetic tweezers assay allows us to probe the folding behaviour of proteins in the presence of different external stimuli. We have performed the experiment with protein L octamer construct, inserted within a C-terminal AviTag and N-terminal HaloTag. The N terminus of the polyprotein is tethered to a glass surface via HaloTag covalent chemistry, while the C terminus is attached with streptavidin-coated paramagnetic bead. Force is applied by introducing a pair of permanent magnets that can exert a magnetic field vertically towards the tethered protein^13,14,17^ (Fig. 1A). Fig. 1B shows a representative trace of protein L, obtained from our single-molecule experiment. At first, a denaturing pulse of 45 pN was applied, which allows the polyprotein to unfold completely to detect the eight unfolding domains of protein L having unfolding extension of 15 nm as a single-molecule fingerprint. Then the force is quenched to 9 pN, allowing the polyprotein to refold. The minimum time takes to unfold all the domains is termed as first passage time (FPT) for unfolding (first inset of Fig. 1B). Similarly, during quenching, the total time required to refold all the polyprotein domains is defined as refolding FPT. Averaging such numerous trajectories, we can determine mean-FPT (MFPT), describing both unfolding and refolding kinetics^16,17,21–23^. Following the complete refolding, the polyprotein hops between folded and unfolded states under force-induced equilibrium condition, which are observed as ascending unfolding steps and descending refolding steps. From this folding-unfolding transition, we estimate the folding probability by dwell time analysis by taking many equilibrium phases from many folding trajectories. In every trajectory, displaying both unfolding and refolding MFPT, each folded domain of the polyprotein is dented as I and their residence time along the equilibrium phase is t=T_I_/T_t_, where T_I_ is the time expend in I state and T_t_ is the observable period of equilibrium phase, characterized as *T*_*t*_ = ∑_*I*_ *T*_*I*_ and N = 8, total number of domains. Thus, FP is calculated as normalized average state (Eq. 1)^15–17,24^,

**Figure 1:**
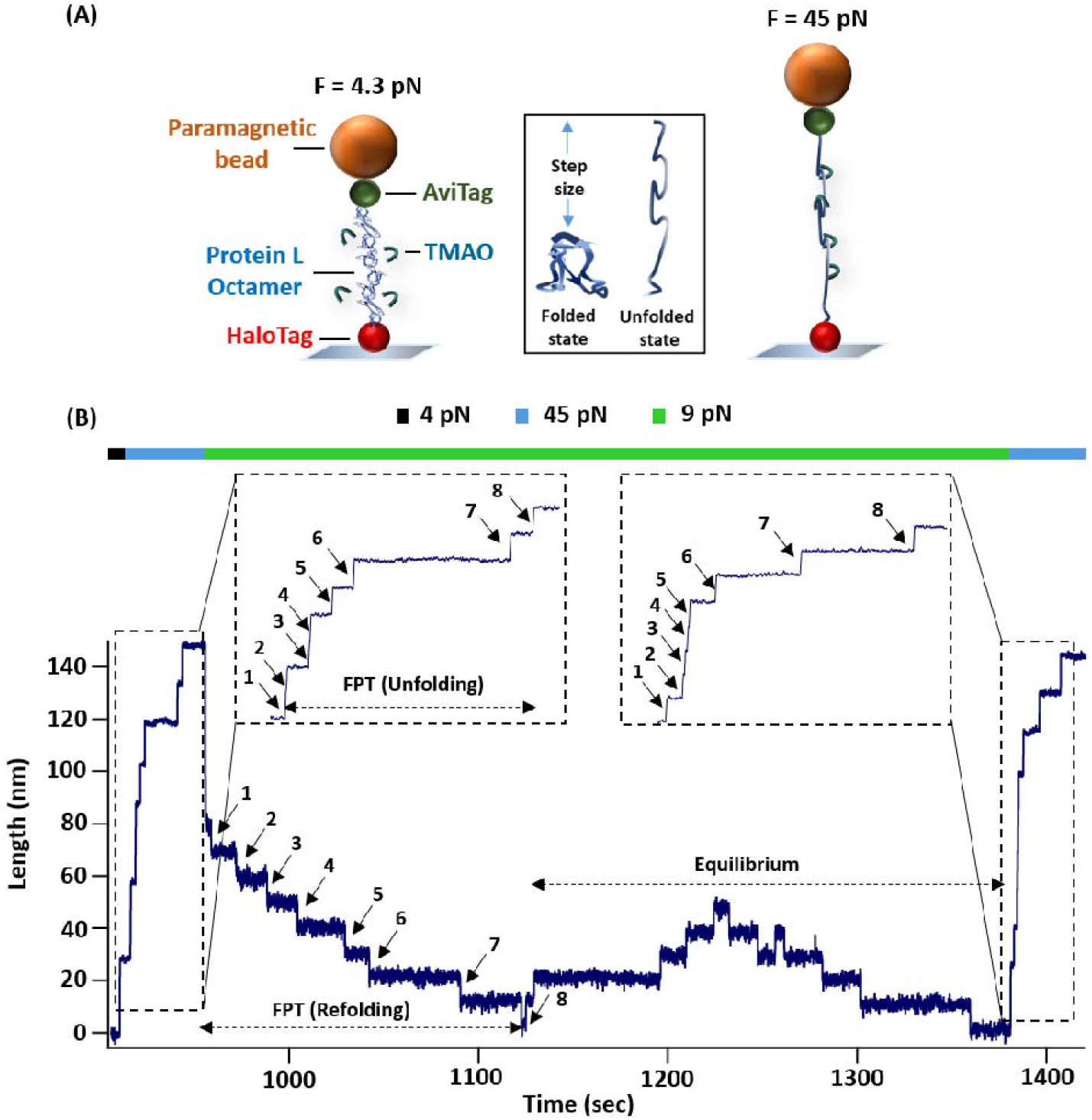
Instrumental setup for magnetic tweezer to study the effect of protein-TMAO interaction: **(A) A schematic illustration of magnetic tweezers experiment:** The polyprotein construct, with eight repeats of protein L domains, is flanked by a C-terminal AviTag for attaching with streptavidin-coated paramagnetic bead and N-terminal HaloTag for tethering with the glass surface. The mechanical force is applied specifically on the protein by neodymium permanent magnet and is controlled by changing its distance with paramagnetic bead. The applied force unfolds the polyprotein, resulting in an extension or step size, which is defined as the difference between unfolded and folded domain length. **(B) Representative trajectory of protein L octamer under force:** At first, the polyprotein construct was fully unfolded at a force of 45 pN, where eight unfolding events were observed as a fingerprint. Then the force was reduced to 9 pN for observing the complete refolding of all protein domains, which is followed by an equilibrium folding transition. To check the unfolding of all the domain, we applied another probe pulse at 45 pN. The total time taken for unfolding of all the domains is defined as first passage time (FPT) for unfolding and similarly, the time taken for complete refolding of all the domains is termed as FPT for refolding.

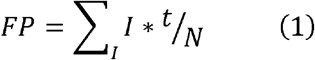

From the folding probability within 8-10 pN force, we also calculated the change in folding free energy difference obtained from the equilibrium condition using Eq. 2.

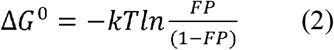

### TMAO increases the mechanical stability of protein L B1

Mechanical strength of protein L has been monitored by force-ramp technology by increasing the force from 4 to 80 pN at a loading rate of 2.53 pN/s. Using protein L octamer construct, we have observed eight distinct unfolding steps while applying the force-ramp protocol (Fig. 2A). The unfolding extensions was vertically aligned with the force-extension curve to estimate the unfolding force of protein L (Fig. 2A, indicated by dotted line).

**Figure 2:**
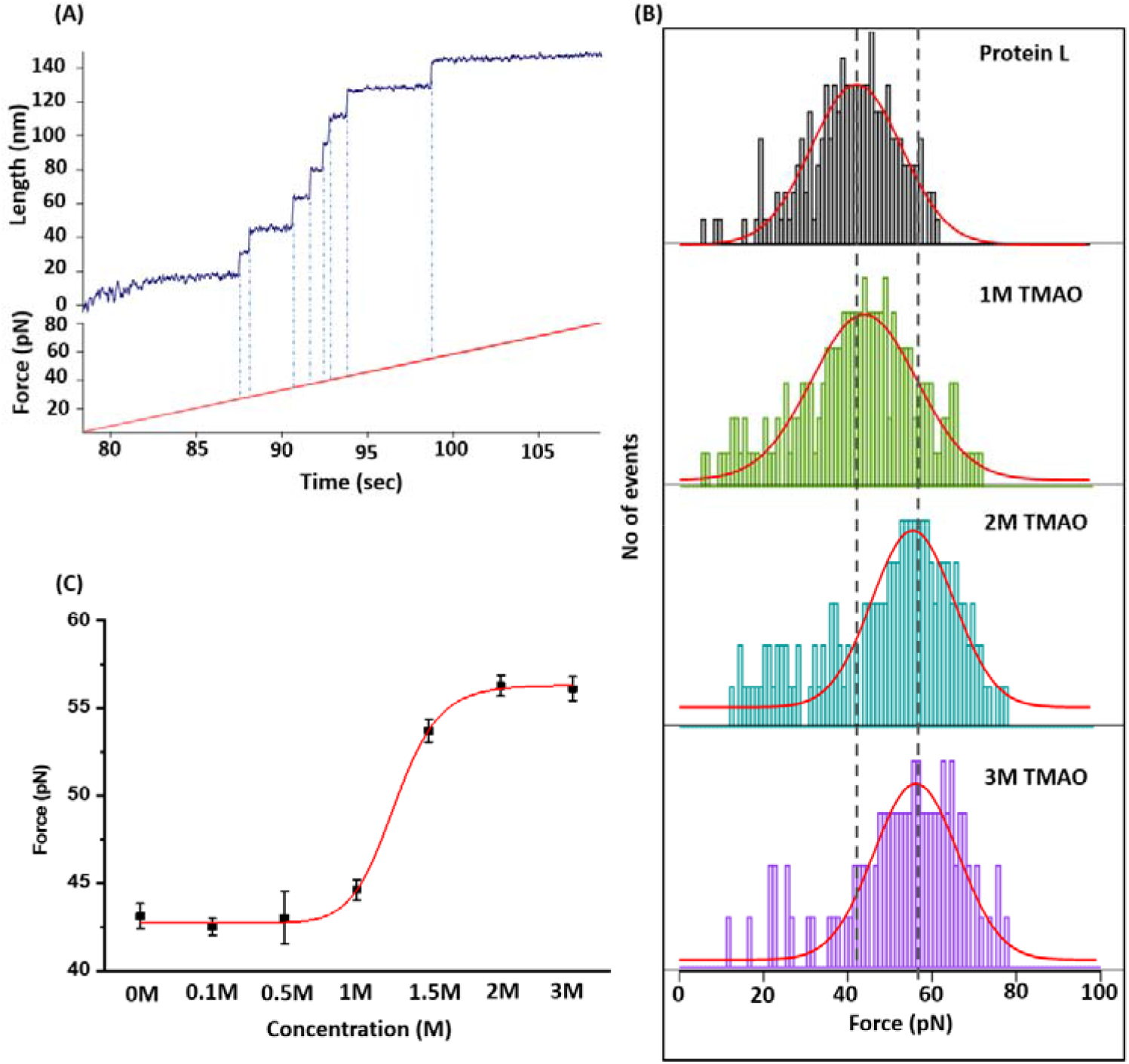
Mechanical strength of protein L modulated by different TMAO concentrations. **(A) Force-ramp experiment of protein L domain:** In the force-ramp protocol, a single protein L octamer is stretched to observe the unfolding force of eight protein L domains at a loading rate of 2.53 pN/sec. **(B) Mechanical stability of protein L in the presence of TMAO:** From the force-ramp experiment, histogram curves with different TMAO concentration have been plotted against the force to monitor its effect on protein L unfolding. Red line represents Gaussian fits to the histogram. As demonstrated, the unfolding force of protein L increases with the TMAO concentrations. **(C) Average unfolding force of protein L:** The average unfolding force, obtained from the histogram plot, are plotted as a function of TMAO concentration. This curve describes that with the increasing TMAO concentration, unfolding force increases in sigmoidal fashion. Error bars are obtained from gaussian fitting of the histogram plot.

We systematically explored TMAO effect on the mechanical stability of protein L by observing their unfolding force using the force-ramp technology. Histogram plots were obtained from these unfolding forces and average unfolding force was measured by fitting the histogram plot with gaussian fit (Fig 1B). Interestingly, we observed that the average unfolding force increases with the TMAO concentrations. For example, the unfolding force is 42.6±0.4 pN without TMAO (control), while increases to 56.2±0.6 pN in the presence of 3M TMAO (Fig. 2B). In Fig. 2C, the average unfolding force has been plotted against the TMAO concentration and observed that the unfolding force changes negligibly upto 1 M TMAO, however, increases significantly upon increasing its concentration to 1.5 M. These data suggest that TMAO significantly increases the mechanical strength of protein L by increasing their average unfolding force, plausibly indicating that TMAO may interact with the folded state of the protein L.

### TMAO modulates the protein L folding events in a concentration-dependent manner

To determine how TMAO affects folding events of protein L, we investigated the unfolding and refolding MFPT of protein L at different TMAO concentrations. Fig. 3A demonstrates the variation of the unfolding MFPT against the force and TMAO concentrations. This plot suggests that at a particular force, the unfolding MFPT increases with the TMAO concentration. For example, at 45 pN, protein L unfolding takes 10.1±1.5 s without TMAO, whereas it increases to 37.3±5.4 s with 3M TMAO. By contrast, refolding MFPT has been observed to decrease in the presence of TMAO (Fig. 3B). These results illustrate that TMAO retards the unfolding kinetics and accelerates the refolding kinetics in concentration-dependent manner.

**Figure 3:**
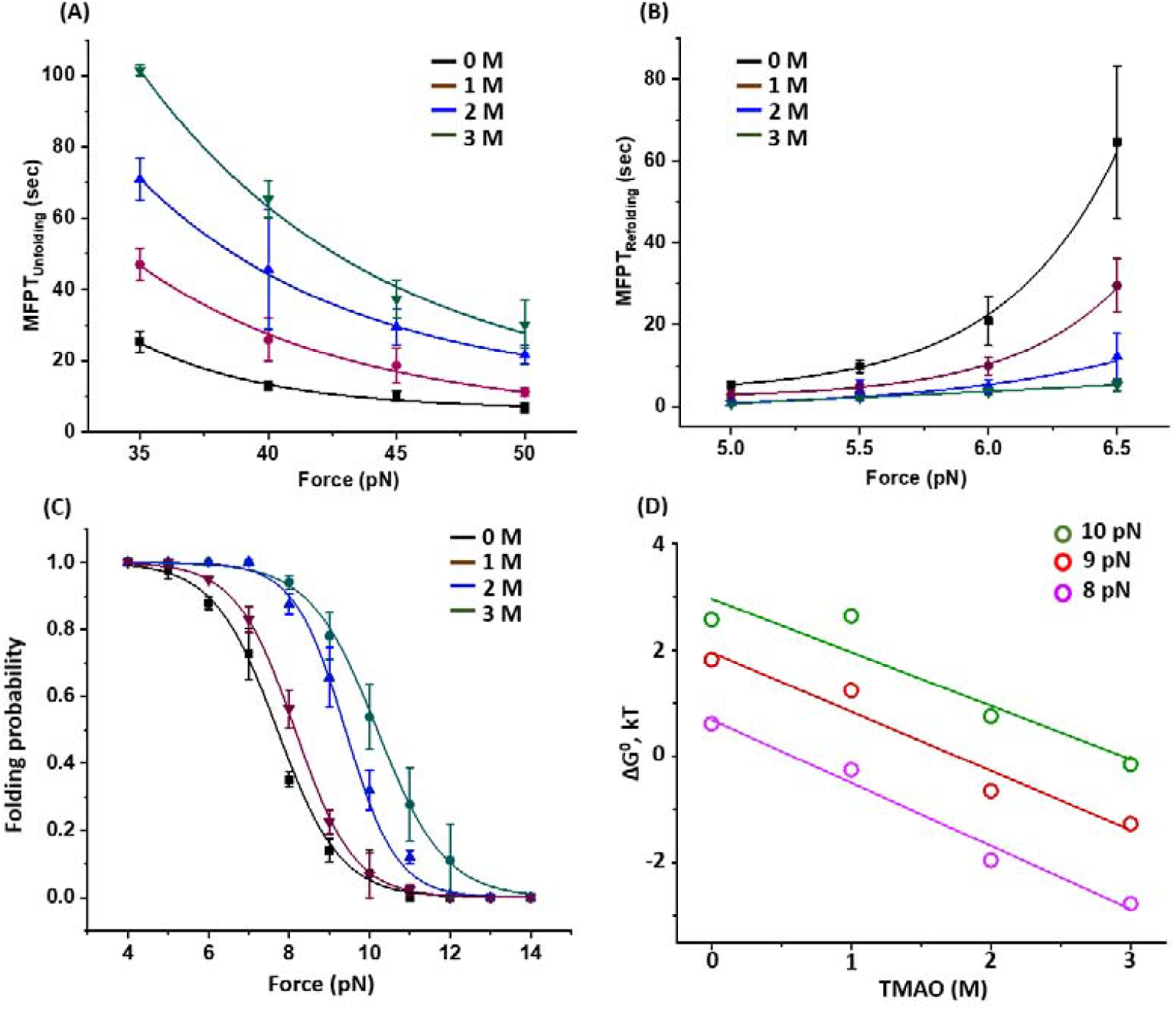
TMAO affects protein L folding events: **(A) Unfolding MFPT of protein L at varying TMAO concentrations:** MFPT unfolding have been plotted at different TMAO concentrations as a function of force. Unfolding time at any particular force increases with the TMAO concentration, signifying that TMAO retards the unfolding kinetics of protein L domain. In each case, more than four individual molecules have been taken for averaging. Error bars are standard error of mean (s.e.m). **(B) Refolding MFPT:** TMAO accelerates the refolding kinetics by decreasing the complete refolding time in a concentration-dependent manner. For example, at 6.5 pN, without TMAO (control), the complete refolding occurs in 64.6±18.7 s, whereas it decreases to 5.34±1.46 s in the presence of 3M TMAO. In each case more than four individual molecules are measured for each force. Error bars are s.e.m. **(C) Comparison between folding probability:** Folding probability of protein L in the presence of different TMAO concentrations have been plotted as a function of force. The effect is mostly pronounced within 7-11 pN range and attains maximal at 9 pN force. It has been observed that TMAO upshifts the half-point force of protein L with the concentration. More than four individual molecules have been taken. Error bars are standard error of mean. **(D) Effect on folding free energy (**Δ**G**^**0**^**) change:** The change in free energy of protein L has been plotted as a function of different concentration of TMAO. From the folding probability data, we calculated the ΔG^0^_Folding_ at different forces within 8 to 10 pN and observed that TMAO decreases the ΔG ^0^_Folding_, signifying that TMAO favors the folding process.

Furthermore, we measured the folding probability (FP) of protein Lin the presence of TMAO over the range of 4 to 14 pN. The drastic change in the folding probability was observed in the intermediate force region of 7-11 pN, with the most pronounced TMAO effect at 9 pN. The half-point force, (the force where FP = 0.5) was increased from 7.7 pN in control to 10.2 pN at 3 M TMAO. This upshift in FP with the TMAO indicates an increased folding ability of protein L under force (Fig. 3C). Additionally, Fig. 3D demonstrates the comparative change in free energy difference within 8-10 pN force, where protein L polyprotein exhibits folding-unfolding dynamics under force-induced equilibrium condition. At 9 pN force, the calculated value *ΔG*^*0*^ of protein L folding is 1.81 kT in the absence of TMAO, whereas it decreases to -1.26 kT at 3 M TMAO. Subsequently, the *ΔΔG*^*0*^ (9 pN) is (*ΔG*^*0*^_*3M*_*-ΔG*^*0*^_*1M*_) is - 3.07 kT, implying the favoured folding process in the presence of TMAO. This result suggests that TMAO have the ability to reshape the free energy landscape by stabilizing the folded state of protein L.

### Mechanical work done by protein L in the presence of TMAO

It is well-established that protein folding generates mechanical energy for performing different cellular processes^24–28^. From our single-molecule data, we observed that TMAO modulates the mechanical work done of protein L folding by tilting the energy landscape towards the folded state. The folding work is measured as the product of the force and step size at that particular force, as described previously^24,25^. This measurement indicates that protein folding at larger force generate more mechanical energy; however, FP values decreases with the force which underestimates the work done calculation. Therefore, a more precise measurement could be performed by multiplying the folding work with the FP under a certain force. Although 3 M TMAO does not affect the protein L step size, it increases the FP data than in its absence; and therefore, a prominent difference has been observed in the work done of protein L folding. We observed that protein L generates a maximum peak of mechanical work at 33.54±5.8 zJ at 6.5 pN (Fig. 4, red circles), whereas 3M TMAO can produce mechanical energy upto 67.3±15.8 zJ at ∼8.3 pN. Thus, TMAO have the ability to delivers ∼2 folds extra mechanical work performed by protein L folding.

**Figure 4:**
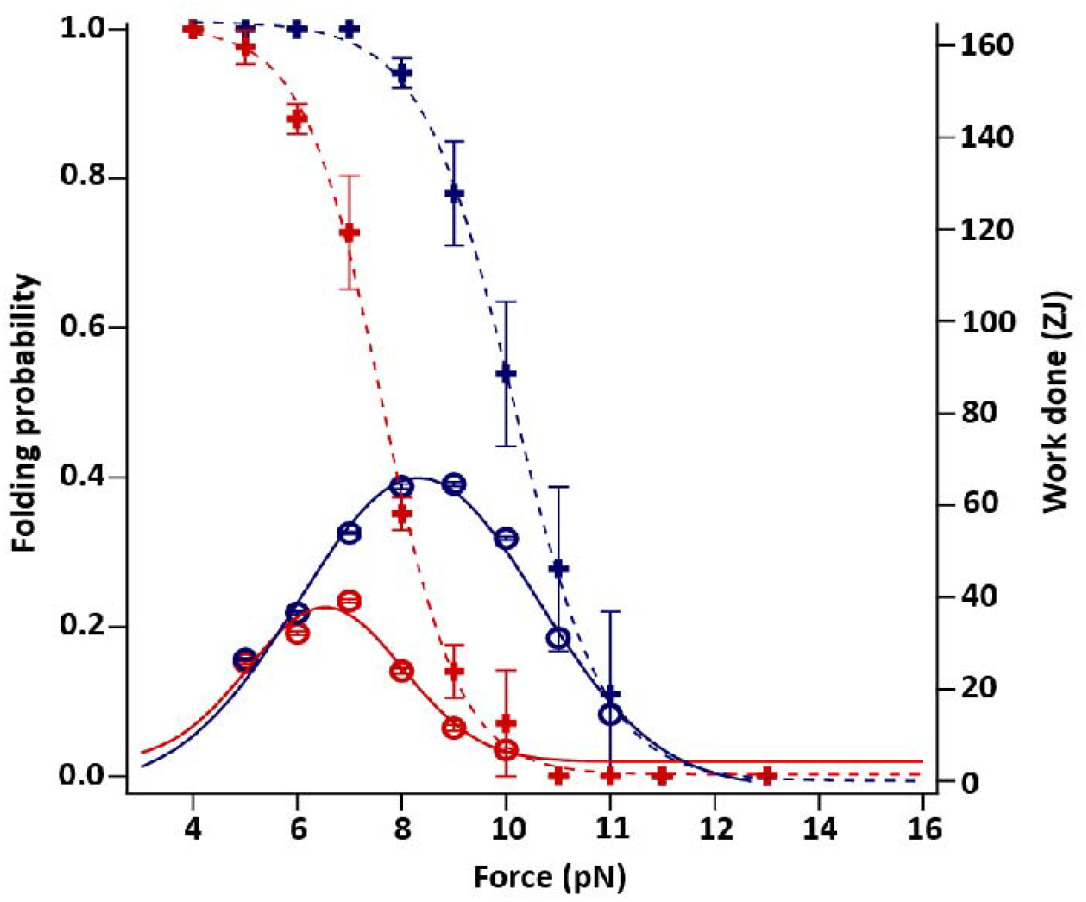
Mechanical work done generated by protein L folding in the presence of TMAO: The value of mechanical work done is calculated as a product of folding work and folding probability, where expected value of folding work is calculated by multiplying step size to force. The highest mechanical energy obtained from gaussian fitting is 33.54±5.8 zJ without TMAO (red circles), while the mechanical energy increases to 67.3±15.8 zJ in the presence of 3M TMAO (blue circles).

## Discussion

Osmolyte molecules are indispensable to prevent urea- and salt-induced denaturation in aquatic animals. TMAO as a protecting osmolyte has been reported to both promote and inhibit the protein folding, though more well-characterized role is assistance in protein folding. Therefore, the precise mechanism of TMAO effect on protein folding is a subject of intense debate. Our single-molecule study reveals that TMAO favours the protein L folding through increasing the protein L unfolding force upto 1.3 folds and thereby, promoting their mechanical stability. Interestingly, we observed a non-significant increase in unfolding force upto 1M TMAO, which is consistent to a previous optical tweezers study on T4 lysozyme^12^. However, the stark changes in the unfolding profile has been observed upon increasing the concentration to 1.5 M, signifying a very high cooperativity in TMAO-protein L binding within a narrow concentration range. Since protein L is functionally nonmechanical, we have also confirmed our study with another mechanosensitive protein talin and observed that TMAO also increases their unfolding force from 12.9 pN to 25.1 pN (Fig. 5A). Despite their structural difference, these two substrates exhibit similar unfolding profile with TMAO, signifying a generic TMAO-mediated mechanical stabilization.

**Figure 5:**
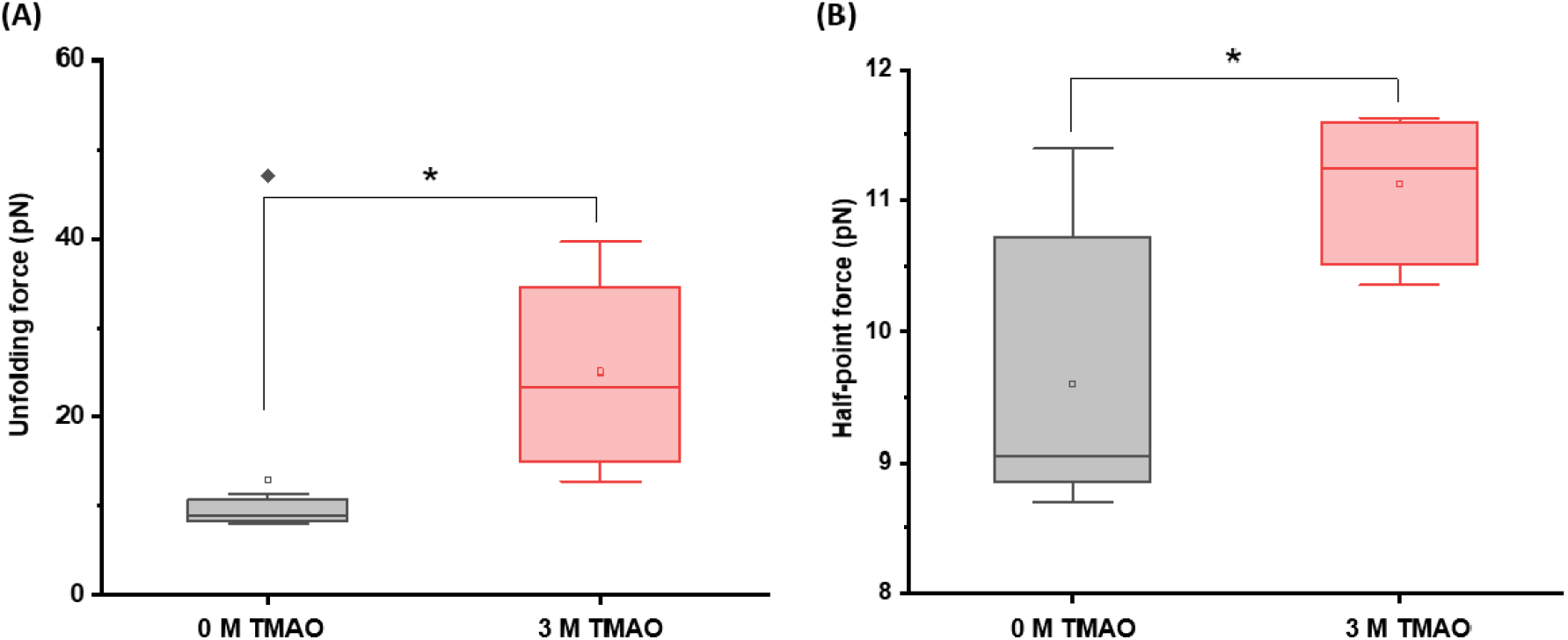
TMAO effect on talin mechanics: **(A) Unfolding force:** TMAO increases the **(A)** unfolding force of talin from 12.9±3.8 pN to 25.1±4 pN and **(B)** half-point force from 9.6±0.4 to 11.1±0.3 pN. Both of these data collectively suggest that TMAO increases the mechanical stability of talin through increasing the unfolding force and half-point force, which in turn helps in the folded state stabilization.

From the kinetics study, it has been observed that TMAO speeds up the refolding kinetics and retards the unfolding kinetics, plausibly stabilizing the folded state of the protein. Since TMAO promotes the native state stabilization of the substrate protein, change in the refolding kinetics was expected. Interestingly, we observed a ∼4.4 folds decrease in the unfolding kinetics, which strongly contradicts the study by Motlagh et al, where TMAO has been found to negligibly change the unfolding kinetics of T4 lysozyme. However, previous single-molecule AFM study revealed that osmolytes in high concentration could slows down the unfolding kinetics and accelerates their compaction under applied tension^29–32^. TMAO effect possibly follows backbone-based osmophobic model, where an osmolyte prevent backbone solvent hydrogen bonding and therefore, destabilizes the unfolded state due to their exposed^33^. This exposure should be compensated by a kinetic protection of mechanical unfolding and faster folding rate^30^. This results in a collapse by decreasing the solubility of peptide backbone due to hydrophobic effect^34^.

Apart from the kinetic study, FP data as well as the free energy calculation also revealed a direct shifting of the folding dynamics towards the folded state. We monitored the folding probability of both the protein L and talin with 3 M TMAO and observed very similar increase in FP (Fig. 5B). Since local TMAO concentration is biologically relevant as orthogonal mechanism to impact on protein folding, this increased FP values in the presence of the TMAO could be important for co-translational folding from the functioning ribosomes. Protein folding at the mouth of ribosomal generate mechanical work output due to confined exit tunnel and scrunching of the translocated polypeptide by the folded portion of the polypeptide. Possibly, TMAO colocalized with the ribosomes could modulate this work output. Indeed, we observed that TMAO can increase the mechanical work done of protein folding almost 2-folds at high concentration. From a concerted view of higher half-point force and mechanical work output revealed that TMAO allows the protein to fold at higher force, which concomitantly generates higher energy than the protein folding in its absence. Overall, this finding concludes that TMAO mechanically stabilizes the substrate protein and therefore, reveals an osmolyte-induced mechanical folding of proteins with single-molecule resolution.

## Author Contributions

S.H designed the project. D.C., A.D., and A.M. performed the experiment. S.H., D.C., S.C., D.C., analysed the data. S.H., S.C., D.C., D.C. wrote the manuscript.

## Acknowledgement

We thank Ashoka University for support and funding. S.H. thanks DBT Ramalingaswami Fellowship and DST SERB Core Research Grant for funding. We sincerely appreciate Prof. Julio Fernandez (Columbia University) for helping us with the magnetic tweezers set up.

